# Targeted insertion in well-characterized *Drosophila* cell lines using φC31 integrase

**DOI:** 10.1101/024216

**Authors:** Lucy Cherbas, Jennifer Hackney, Lei Gong, Claire Salzer, Eric Mauser, Dayu Zhang, Peter Cherbas

**Author notes:** Corresponding author: Lucy Cherbas Department of Biology Jordan Hall 1001 East Third Street Bloomington IN 47405. current address: School of Mathematics and Natural Science, New College, Arizona State University, Phoenix AZ. current address: Hangzhou School of Agriculture and Food Science, Zhejiang Agriculture and Forestry University, Zhejiang Lin’an, Hangzhou, China (Dayu Zhang).

## Abstract

We describe an adaptation of φC31 integrase-mediated targeted cassette exchange for use in *Drosophila* cell lines. Single copies of an attP-bounded docking platform carrying a GFP-expression marker, with and without insulator elements flanking the attP sites, were inserted by P-element transformation into the Kc167 and Sg4 cell lines; each of the resulting docking site lines carries a single mapped copy of one of the docking platforms. Vectors for targeted substitution contain a cloning cassette flanked by attB sites. Targeted substitution occurs by integrase-mediated substitution between the attP sites (integrated) and the attB sites (vector). We describe procedures for isolating cells carrying the substitutions and for eliminating the products of secondary off-target events. We demonstrated the technology by integrating a cassette containing a Cu^++^-inducible mCherry marker, and we report the expression properties of those lines. When compared with clonal lines made by traditional transformation methods, which lead to the illegitimate insertion of tandem arrays, targeted insertion lines give more uniform expression, lower basal expression and higher induction ratios. Targeted substitution, though intricate, affords results that should greatly improve comparative expression assays – a major emphasis of cell-based studies.

## INTRODUCTION

Stable cell lines have formed an increasingly useful portion of the *Drosophila melanogaster* tool kit in recent years, as the number of readily available lines has rapidly expanded, and many of those lines have been characterized extensively (CHERBAS AND GONG 2014). Over 100 diverse lines came are now available through a cell line stock center maintained by the Drosophila Genomics Resource Center (DGRC); molecular characterization of many of the lines has occured in many laboratories, both as part of the modENCODE project and independently (Zurovec *et al.* 2002; Dasgupta *et al.* 2005; Williams *et al.* 2007; Lau *et al.* 2009; Liu *et al.* 2009; Schaaf *et al.* 2009; Schwartz *et al.* 2010; Cherbas *et al.* 2011; Eaton *et al.* 2011; Koppen *et al.* 2011; Riddle *et al.* 2011; Vatolina *et al.* 2011; Alekseyenko *et al.* 2012; Riddle *et al.* 2012; Brown *et al.* 2014; Lee *et al.* 2014b; Wen *et al.* 2014).

Stable transformation is a widely used tool in both flies and their cell lines; its power has increased in recent years as the random insertions of P elements has been supplemented by site-directed insertions of DNA into the chromosomes of flies. The use of integrase from the bacteriophage φC31 to perform site-specific recombination is a particularly popular version of the latter approach (Huang *et al.* 2009a; Ejsmont and Hassan 2014). Originally developed for mammalian systems, this technique is now well established in flies (Groth *et al.* 2004; Venken *et al.* 2006; Fish *et al.* 2007; Huang *et al.* 2009a; Venken and Bellen 2012); it has been used for simple insertion of plasmids and much larger constructs (Venken *et al.* 2010) through the recombination of a single attP site (either pre-existing in the genome or inserted into the chromosome) with a single attB site in the targeting construct. It has also been used to mediate cassette exchange, in which a chromosomal DNA sequence bounded by attP sites is exchanged for a plasmid sequence bounded by attB sites (Bateman *et al.* 2006; Fujioka *et al.* 2008; Huang *et al.* 2009b; Weng *et al.* 2009; Bateman *et al.* 2012; Sun *et al.* 2012; Bateman *et al.* 2013; Zhang *et al.* 2014). The integrase is produced either from injected RNA (Groth *et al.* 2004; Fish *et al.* 2007) or from a stably integrated φC31 integrase transcription unit which can be removed in a subsequent genetic cross (Bischof *et al.* 2007). Targeted insertions and cassette exchanges make possible the repeated integration of constructs into an identical DNA environment, thereby eliminating the variations caused by position effects.

In cell lines, φC31 integrase-mediated targeting would confer improvements to currently used techniques beyond those seen in flies. Current techniques for stable transformation of *Drosophila* cell lines stably transformed by techniques now in common use lead to the formation of tandem arrays of the transforming plasmid, often quite long, which are inserted by illegitimate recombination into the genome(Bourouis and Jarry 1983; Moss 1985; Cherbas *et al.* 1994b). This anomalous structure, which is also seen in transformed cells from mammals (Wurtele *et al.* 2003; Rosser and An 2010) and to an extreme degree in a mosquito cell line (Monroe *et al.* 1992), leads to abnormal chromatin structure, silencing of expression (Rosser and An 2010), pairing between arrays (Mirkin *et al.* 2014), abnormal regulation caused by saturation of the supply of critical *cis*-acting factors, and instability in the length of the array. The resulting effects on regulation of transgene expression and the cell-to-cell variability in transformed lines, even after cloning, provide strong incentives to adapt targeted transformation techniques for cell lines. φC31 integrase-mediated gene targeting has been used in mammalian cell lines (Goetze *et al.* 2005), and the integrase has been shown to function in the *Drosophila* cell line S2 (Groth *et al.* 2004). But targeted integration in cell lines has proved somewhat difficult, and to our knowledge, the system has been pursued in only three laboratories: The Perrimon laboratory placed MiMIC elements, an enhancer-trap version of a φC31 docking site, into S2R+ cells, and briefly described an integrase-mediated cassette exchange as a proof-of-principle (Neumuller *et al.* 2012). The Simcox laboratory has used the alternative approach of making new cell lines from flies carrying well-characterized attP docking platforms (Manivannan *et al.* 2015). In our laboratory, as described in this paper, we have placed single copies of φC31 docking platforms into well-characterized pre-existing cell lines using P element transformation of the cell lines, and established conditions for carrying out φC31 integrase-mediated exchange at these docking platforms. Here, we describe the generation of a set of tools for targeted insertion of constructs into the *Drosophila* cell lines Kc167 and Sg4. We will describe in detail cassette exchange in two of the Kc167 docking site lines, and compare the properties of the products of targeted exchange with those of stably transformed lines made with the same transgenes by more traditional means.

## METHODS

### Cell culture

Kc167 and Sg4 cells were obtained from the Drosophila Genomics Resource Center; the former is a clone of Kc (Echalier and Ohanessian 1969; Bourouis and Jarry 1983), the latter a clone of S2 (Schneider 1972; Arndt-Jovin 2006). Kc167 cells were grown in the serum-free medium CCM-3 (GE Healthcare HyClone) unless otherwise indicated; Sg4 were grown in Shields and Sang medium M3 with added bactopeptone and yeast extract (M3+BPYE (Cherbas *et al.* 1994b)) supplemented with 10% heat-inactivated fetal calf serum. General procedures for cell culture were as described previously (Cherbas *et al.* 1994b).

Cells were cloned by a modification of a procedure described previously (Cherbas *et al.* 1994b; Cherbas and Cherbas 2007). A feeder layer was prepared from cells of the parental line (Kc167 or Sg4) by pelleting cells, resuspending them in 5 ml Robb’s saline (Robb 1969) in a 25 cm^2^ T flask, and exposing them to 60 kR of gamma rays in a cesium source. The irradiated cells were then transferred to M3+BPYE supplemented with penicillin (100 u/ml), streptomycin (100 μg/ml), and heat-treated fetal calf serum (5% for Kc167, 10% for Sg4), at a final concentration of 1.5 × 10^6^ cells/ml. This feeder cell suspension was plated in 96-well plates, at 100 μl/well. Cells to be cloned were dispensed individually into the wells, using a fluorescence-activated cell sorter (see below). After approximately 2 weeks, clones were picked, scaled up in their normal medium (CCM-3 for Kc167, M3+BPYE + 10% serum for Sg4) as described (Cherbas *et al.* 1994b), and used for analysis and for the preparation of frozen stocks. Cloning efficiency was typically 10-20%. Although Kc167 cells and their derivatives were normally maintained in CCM-3 medium, their cloning efficiency was near zero if they were dispensed by the cell-sorter into a feeder-cell suspension in CCM-3; for that reason we used M3+BPYE with 5% serum for the feeder-cell suspension, and reverted to CCM-3 for expansion of the growing clones.

### Plasmid construction

Sequences for all of the plasmids constructed for the experiments described in this paper are available in Supplemental Materials. We constructed two types of docking sites in the P element vector Carnegie4 (Rubin and Spradling 1983b), with and without *gypsy* insulator elements flanking a pair of parallel φC31 attP sites. Both docking sites contain a nuclear eGFP expression cassette between the attP sites. Maps for the 2 docking site transposons are shown in Figure 1A. Vectors for targeting to the docking sites are shown in Figure 1B; these plasmids each contain a pair of parallel φC31 attB sites flanking a methotrexate resistance marker, with a herpes simplex TK expression cassette conferring ganciclivir sensitivity located outside the attB sites. One of the vectors also contains a Gateway insertion cassette for use in inserting fragments to be transported to the docking site. Figure 1C shows the attB-bounded region containing Mt-mCherry inserted into the Gateway site for use in targeting these constructs to docking sites. In act-ϕC31-integrase, the coding sequence for φC31-integrase was placed under the control of a strong constitutive promoter from *Act5C*. Complete sequences for all of these plasmids are given in the supplemental materials; critical portions came from the following plasmids: insulators and eGFP from pStinger (Barolo *et al.* 2000), P element ends from Carnegie4 (Rubin and Spradling 1983a), actin promoter and DHFR coding sequence from pUC-act-DHFR (Segal *et al.* 1996), metallothionein promoter from pRmHa-1 (Bunch *et al.* 1988), ϕC31 integrase coding sequence from pET11phiC31polyA (Groth *et al.* 2004), HSV TK coding sequence from pAL119-TK (Dewey *et al.* 1999) (from AddGene), attP sites from pXLBacII-attP-yellow forward (gift from Koen Venken), attB sites from attB-P[acman]-ApR (Venken *et al.* 2006), Gateway entry cassette (from Invitrogen), mCherry coding sequence from pmCherry Vector (from Clontech). Except where otherwise indicated, all of the source plasmids were obtained from the vector collection of the Drosophila Genomics Resource Center.

**Figure 1.**
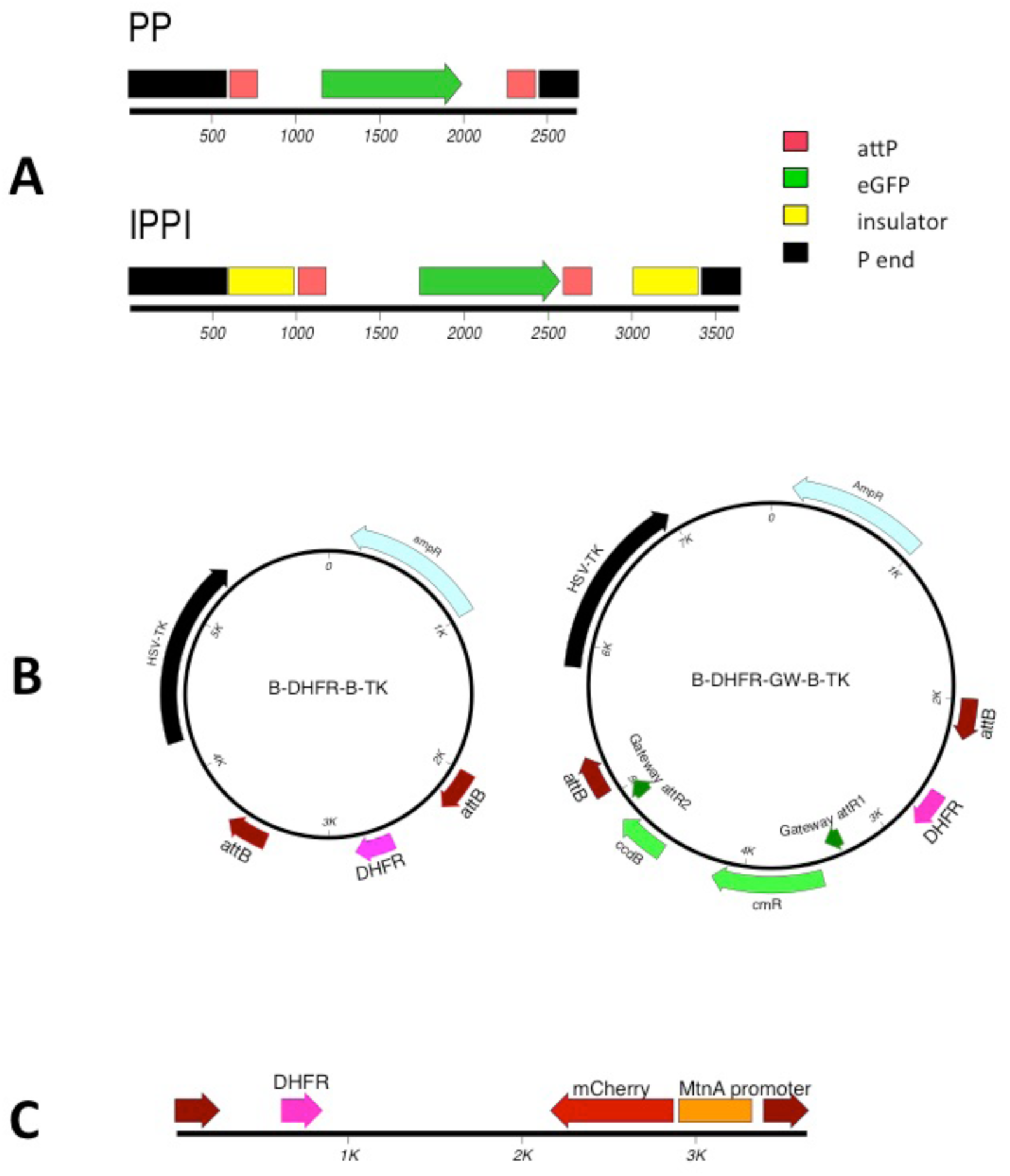
Constructs used in this paper. Sequences for these plasmids and the integrase expression plasmid are provided in the supplemental materials. **A**. The P transposons used as docking platforms. Each contains a GFP expression cassette between parallel attP sites; IPPI also contains insulator elements flanking the attP sites. Only the P element transposons are shown; see supplemental materials for the sequence of the whole plasmids used for transposition of these docking platforms. **B**. Vectors for integration by replacement in the docking sites. Each contains a methotrexate-resistance marker between parallel attB sites for positive selection and a HSV-TK expression cassette outside the attB-bounded region, for counter-selection against cells that have acquired the entire plasmid by illegitimate integration. B-DHFR-GW-B-TK has a cassette for inserting constructs using the Gateway system (Life Technologies, Inc.). B-DHFR-B-TK has a limited number of unique sites for inserting constructs, including an EcRI site upstream of the DHFR transcription unit and a ClaI site downstream of DHFR. **C.** A Cu^++^-inducible mCherry targeting sequence used for integration into the docking sites in experiments described in this paper. The targeting plasmid was made by cloning an Mt-mCherry transcription unit into the Gateway entry vector pENTRB (Life Technologies), and then using the Gateway reaction to place the transcription unit into B-DHFR-GW-B-TK (shown in Panel B). Only the fragment bounded by attB sites is illustrated here; see Supplemental materials for the sequence of the complete targeting plasmid.

### Microscopy

To screen clones for expression of GFP, we examined clones growing in the original 96-well plates into which they had been sorted, using a BD Pathway 435 High-Content Bioimager. For photomicrography, we placed 1 ml of growing cells into a 35 mm petri dish with a poly-lysine-coated glass bottom (MatTek Corp.); after the cells had settled onto the surface, they were visualized using an Applied Precision DeltaVision personal DV Live Cell Imaging System. Both of these microscopes are housed in the Light Microscopy Imaging Center of Indiana University, Bloomington.

### Fluorescence-activated cell sorting

All cell sorting and cloning were carried out in the Flow Cytometry Core Facility of Indiana University, Bloomington. Populations for sorting were selected for single cells by light scatter, and for living cells by either light scatter (FSC-A vs. SSC-A) or by exclusion of propidium iodide dye. eGFP was excited with a 488 nm 100 mW or 30 mW laser, with emission detected with a 530/30 bandpass filter. mCherry was excited by a 561 nm laser at 150 mW, with emission detected at 610/10. Propidium iodide was excited with a 561 nm 150 mW laser and detected at 582/15. Cloning, with or without fluorescence selection, was done on a FACSAria IIu machine (BD Biosciences). Analysis without cloning was carried out on either the FACSAria IIu or an LSRII flow cytometer (BD Biosciences).

### Molecular analysis by PCR

For digital drop PCR (ddPCR), DNA was prepared from approximately 1.5 × 10^6^ cells using a QIAamp DNA Micro Kit (Qiagen), yielding a final volume of 30 μL. For all other forms of PCR, we used a cell lysate (Gloor *et al.* 1993), modified as follows: Approximately 1.5 × 10^6^ cells were centrifuged and the culture medium removed. Pelleted cells were resuspended in 50μL of squishing buffer (SB: 10mM Tris-Cl pH8.2, 1mM EDTA, 25mM NaCl and 200μg/ml Proteinase K (Qiagen)), and incubated at 37°C for 30 min. The lysate was then heated to 95°C for 2 minutes to inactivate Proteinase K.

Copy number was determined by digital droplet PCR. 8.5 μL of DNA was digested with 10 units of EcoRI-HF (New England Biolabs) for 1 hr at 37°C. Following a 20 minute incubation at 65° to inactivate the enzyme, 1uL of the digest was used for each 20uL ddPCR reaction. Primer sequences are given in the supplementary materials. All copy number variation assays were duplexed with an *EcR* reference assay; the *EcR* region is known to be present in 4 copies in Kc cells (Cherbas and Cherbas 1997; Lee *et al.* 2014a). Reactions were set up using 2x ddPCR Super Mix for Probes (Bio-Rad), each 20x primer and probes (CNV assay and reference assay), and digested DNA in a final volume of 20uL. Digital droplet PCR (BioRad) was set up and performed as described Hindson *et al.* (2011). Thermal cycling conditions for reaction emulsifications (Eppendorf Mastercycler) were 95° for 1 min, 94° for 30s and 62° for 30s (40x, 50% ramp speed), 98° for 10 min followed by a 4° hold.

For clones which had a single copy of a docking site, the insertion site of the docking site transposon was mapped by splinkerette PCR (Potter and Luo 2010), using 25uL of cell lysate in place of the purified genomic DNA in the original protocol.

We used conventional PCR for additional characterization of docking sites and targeted insertion. Sequences of primers used for all PCR techniques are given in the Supplemental Materials.

### Transfections for integrase-mediated insertion into the docking sites

Docking site lines were transfected using Lipofectamine LTX with PLUS reagent (Life Technologies) and a mixture of the integrase-expression plasmid and an attB-targeting plasmid. according to the manufacturer’s protocol. Four days after transfection, we began selection with methotrexate (MTX) as described previously (Cherbas *et al.* 1994b). After the MTX-sensitive cells had died and been replaced with a MTX-resistant population (approximately 2 weeks), we added ganciclovir (GCV), while continuing the MTX selection. Approximately 2 weeks later, we removed the selective agents and cloned GFP^-^ cells.

### Reagent and data availability

All cell lines and plasmids described in this paper are available from the Drosophila Genomics Resource Center (https://dgrc.bio.indiana.edu). Files S1 and S2 contain sequences of plasmids generated in this work and of PCR primers, respectively.

## RESULTS

### Insertion of docking sites into the genome of *Drosophila* cell lines.

Commonly used methods of transforming *Drosophila* cells generate multiple copies in tandem arrays. In order to insert single copies of a docking site into cultured cells, we used P element transposition, as described previously (Segal *et al.* 1996). We began with Kc167 cells, which were used in the earlier work on P element transposition in cells, and subsequently repeated the procedure with the S2 derivative Sg4. In both cases, the transfection efficiency was low (as expected with electroporation), and in order to clone stably transformed GFP-expressing cells, we found it necessary to include an intermediate sorting step. We collected GFP-expressing cells 4 days after transfection, a time when much of the expression is still coming from plasmids not stably incorporated into the genome, to generate a population enriched for transformed cells. GFP-positive cells were cloned from the enriched population approximately 7-10 days later. Once the clones were large enough to visualize, we screened them for GFP expression in a fluorescence microscope. This last step was used to eliminate roughly 30% of the clones, which we presume derived from cells that either were transiently expressing GFP at the time of cloning and or whose autofluorescence caused them to be scored as GFP^+^ by the FACS. Autofluorescence, a significant source of error in FACS because the range of GFP fluorescence from cells carrying a single copy of the transposon overlaps the range of autofluorescence, is easily easily distinguished from GFP fluorescence in microscopy because autofluorescence is punctate and cytoplasmic while GFP expression in these cells is nuclear.

The number of copies of the docking site was determined by ddPCR for each clone, and clones carrying a single docking site per cell (6-40% of GFP-expressing clones in 3 experiments with Kc167, 20-30% in 2 experiments with Sg4) were expanded, saved as frozen stocks and used for further analysis. For each single-copy clone, the insertion site of the docking site was determined by splinkerette PCR (Potter and Luo 2010) and duplicate clones were discarded. Mapping of the insertion sites was confirmed by PCR, using primers from the genomic regions flanking the insertion site. Table 1 lists the docking site clones that we recovered: One IPPI insertion and 10 PP insertions in Kc167, and 6 PP insertions in Sg4.

**Table 1.** Positions of docking site insertions in *Drosophila* cell lines. Names of lines are in the format [parental line] - [type of docking site (PP or IPPI)] – [site of insertion (given as the polytene region containing the insertion site)]. Molecular coordinates refer to the *Drosophila* melanogaster genome, release 6. In those cases where the coordinate is not given precisely, our sequencing reached within a few bases of the recombination site, but did not cross the junction between the docking site and the chromosomal sequence. The direction of the insert is shown with 5’ taken as the left end of the map shown in Figure 1A.

| Name of line | Molecular coordinate of insertion | Direction of insertion | Nearest annotated gene |
| --- | --- | --- | --- |
| Kc167-PP-16F | X:18,094,755 | 5' towards centromere | *RhoGAP16F* |
| Kc167-PP-21B | 2L:161,526 | 5' towards centromere | *spen* |
| Kc167-PP-21D | 2L:479,848 | 5' towards centromere | *cbt* |
| Kc167-PP-50Aa | 2R:13,343,299 | 5' towards telomere | *CR44206* |
| Kc167-PP-50Ab | 2R:13,337,584 | 5' towards centromere | *CR44206* |
| Kc167-PP-52E | 2R:16,125,298 | 5' towards telomere | *spin* |
| Kc167-PP-61C | 3L:635,370 | 5' towards telomere | *CR43334* |
| Kc167-PP-89B | 3R:>=16,189,680 | 5' towards telomere | *sra* |
| Kc167-PP-93E | 3R:<21,591,314 | 5' towards telomere | *InR* |
| Kc167-PP-99A | 3R:29,287,394 | 5' towards telomere | *CG14506* (10 kb away) |
| Sg4-PP-3A | X:2,545,583 | 5' towards centromere | *trol* |
| Sg4-PP-27F | 2L:7,423,926 | 5' towards centromere | *CR43857* |
| Sg4-PP-49B | 2R:12,589,942 | 5' towards telomere | *Sin3A* |
| Sg4-PP-57B | 2R:20,961,063 | 5' towards centromere | *hbn* |
| Sg4-PP-70F | 3L:14,757,988 | 5' towards centromere | *Trl* |
| Sg4-PP-84E | 3R:8,106,183 | 5' towards telomere | *puc* |
| Kc167-IPPI-66D | 3L:8,686,703 | 5' towards telomere | *h* (7 kb away) |

Each docking site clone has significant variation in the intensity of the GFP signal of individual cells, and includes a small fraction of cells in which no GFP expression is detected either by FACS or by fluorescence microscopy (Figure 3). In Kc167-PP-93E, a typical docking site line, approximately 1% of the population has no detectable GFP fluorescence. To characterize the GFP-null cells in Kc167-PP-93E we stained cells with Hoechst 33342 and analyzed by FACS to estimate their position in the cell cycle. There was no significant difference in the distribution of Hoechst 33342 staining between the population as a whole and the GFP-null portion of the population (data not shown); hence loss of GFP fluorescence does not appear to be associated with a stage of the cell cycle. We cloned separately GFP-expressing and GFP-null subpopulations of Kc167-PP-93E. GFP-expressing cells gave rise to healthy clones, each of which had a subpopulation of GFP-null cells indistinguishable from that of parental population (1.16 ± 0.21% for 4 clones). By contrast, GFP-null cells gave rise to small, unhealthy-looking clones composed overwhelmingly of GFP^-^ cells. These observations suggest that occasionally Kc167-PP-93E cells permanently lose their ability to express GFP, but those cells grow poorly, leading to a steady-state level of a few percent. We speculate that loss of GFO expression occurs when a chromosomal rearrangement removes all or part of the docking site. The only docking-site lines in which significantly more than 1-2% of cells fail to express GFP are two lines in which the transposon is inserted near the tip of chromosome arm 2L (Kc167-PP-21B and -21D; see Figure 2); perhaps loss of one copy of this region may is either more frequent or less deleterious than loss of other regions in which docking sites have inserted.

**Figure 2.**
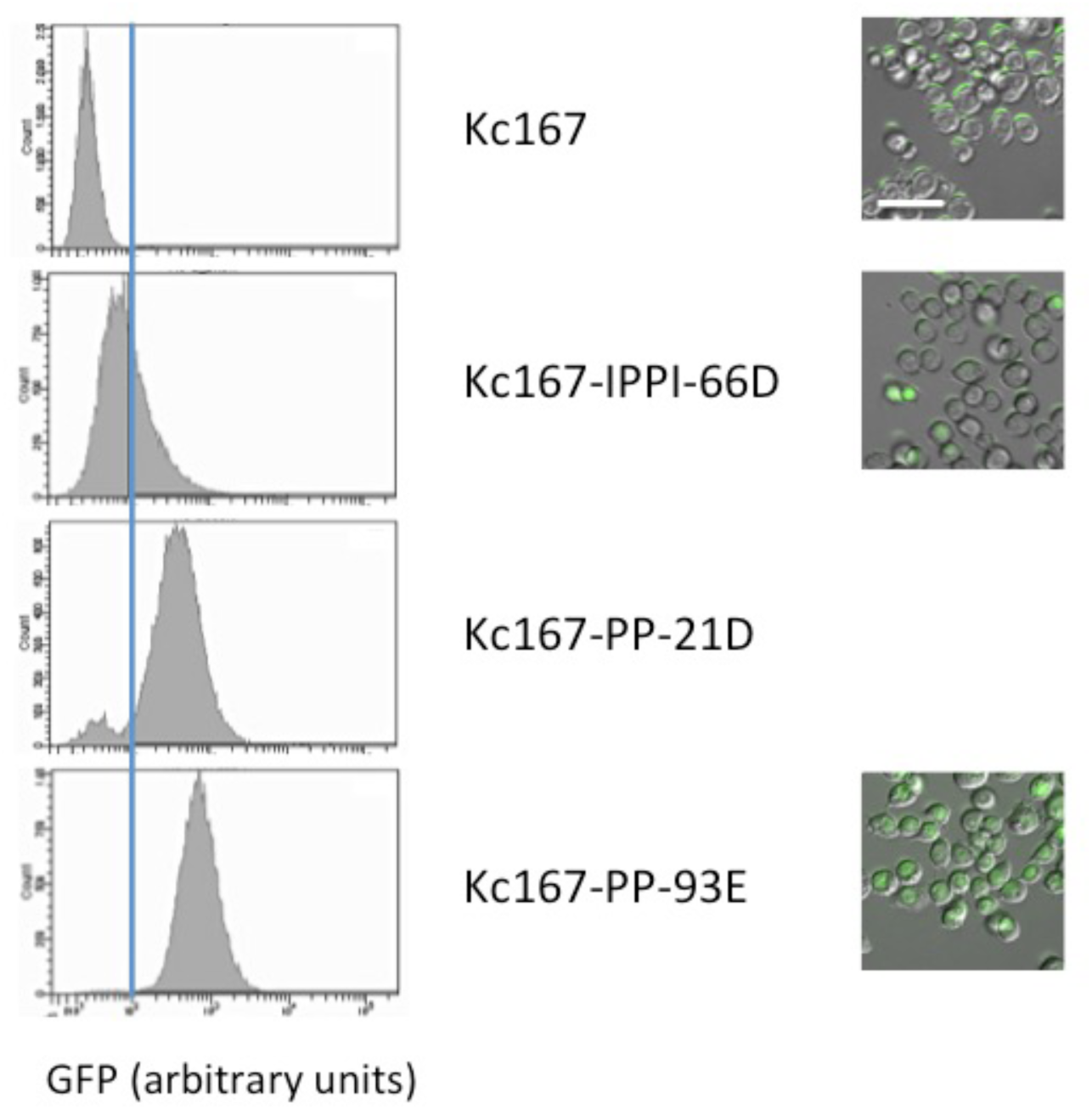
Expression of GFP in docking site lines. FACS-generated histograms are shown at left; fluorescence photomicrographs, at right. Kc167 is the untransformed parental line, exhibiting only autofluorescence. Representative clonal docking site lines are shown below. The vertical blue line at 100 units GFP is provided for visual alignment. GFP-null cells are estimated at about 10% of the population in Kc167-PP-21D and 1% in Kc167-PP-93E. Any GFP-null cells in Kc167-IPPI-66D are masked by the overlapping range of GFP expression. The scale bar for the micrographs represents 25 μm.

**Figure 3.**
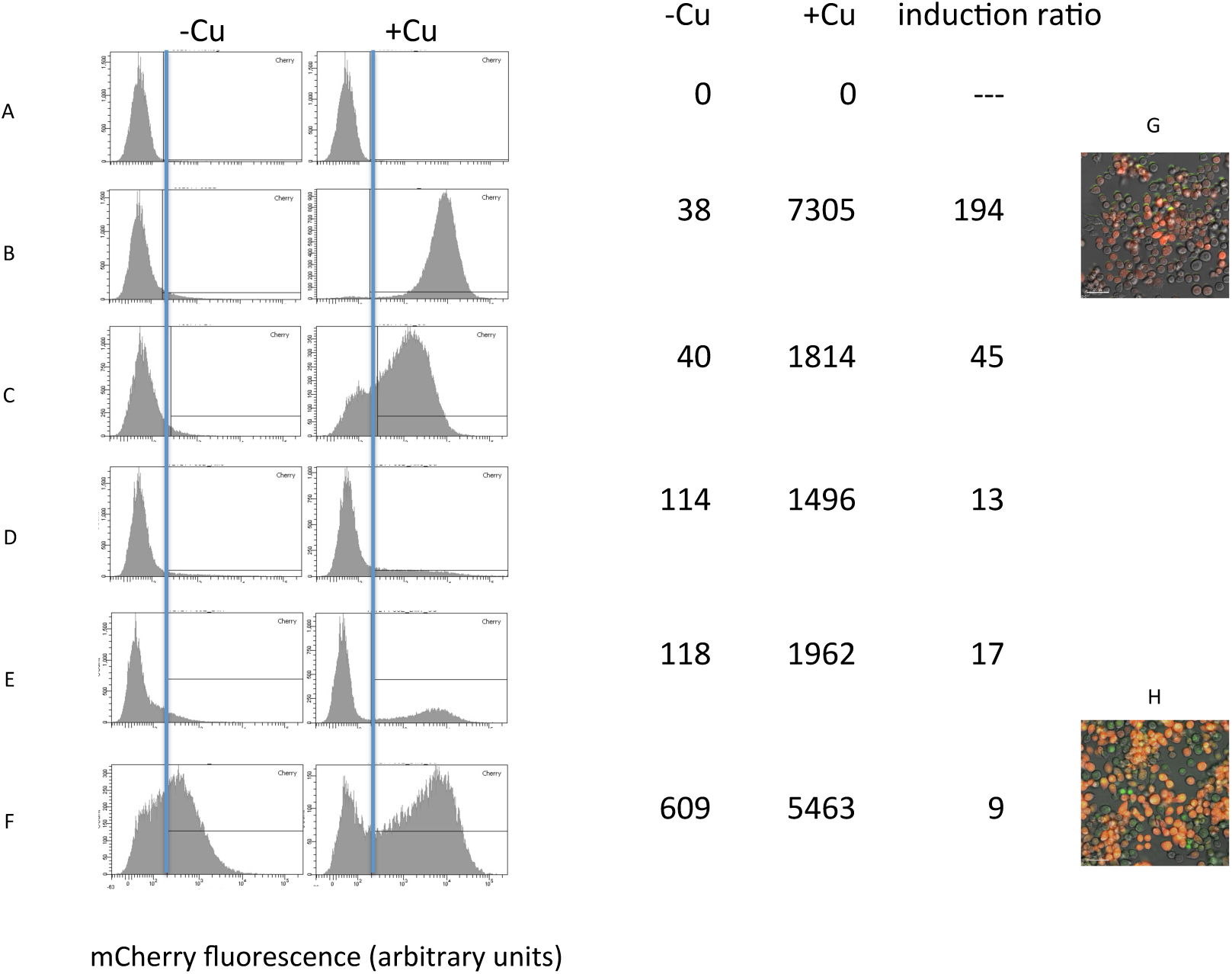
Expression of a transgene in transformed clones. mCherry expression (arbitrary units) was measured by FACS analysis, and is shown as histograms of cell populations with or without Cu++ treatment (1 mM CuSO_4_, 20 hr), and as mean expression and Cu^++^ induction ratio. A: Kc167 (parental line); these cells do not contain an mCherry coding sequence; background autofluorescence measured in these cells is subtracted to give the mean mCherry expression estimates shown in the –Cu and +Cu columns. B, C: targeted substitution of Mt-mCherry into the docking site lines Kc167-IPPI-66D and Kc167-PP-93E, respectively. D, E, and F: illegitimate insertions of the Mt-mCherry plasmid, derived from transformation of Kc167-IPPI-66D (D) or Kc167-PP-93E (E and F) in the absence of φC31 integrase, followed by MTX selection and cloning G, H: fluorescence photomicrographs of targeted substitution of Mt-mCherry in Kc167-IPPI-66D (G) and illegitimate transformation of the docking site line Kc167-PP-93E (H); both cultures were treated with CuSO_4_ to induce mCherry expression. Note the loss of nuclear GFP in panel G, and its retention in panel H.

The cell-to-cell variation of GFP expression is easily visible in fluorescence microscopy (Figure 2; note that the FACS and imaging data are not directly comparable, because the linearity properties of the two techniques are quite different). The basis for this variability is unknown, but we observed a similar level of variation in another cell line which has a single copy of an unrelated GFP transgene: Jupiter (Karpova *et al.* 2006) (data not shown.

### Targeted insertion by cassette exchange.

We describe here the insertion of targeting constructs into two docking site lines: Kc167-IPPI-66D and Kc167-PP-93E. We expect that similar procedures will give successful insertions in the remaining Kc167 and Sg4 docking site lines.

When cells containing a docking platform marked by GFP expression were challenged with a plasmid which contained a methotrexate-resistance marker between two attB sites, in combination with an integrase-expressing plasmid, correctly integrated products would be resistant to methotrexate and would fail to express GFP; we initially tried to select targeted integration using either or both of these properties. Two problems, described individually below, necessitated modifications to this scheme.

1. Loss of GFP was usually associated with loss of the entire docking platform, rather than with targeted substitution. This became obvious when we cloned GFP^-^ cells, and examined the DNA of individual clones, using PCR primers flanking targeted to genomic sequences flanking the docking platform insertion site. Both Kc167 and Sg4 are tetraploid at most loci (LEE *et al.* 2014b); hence PCR reaction of each docking site line produces a small amplicon from the 3 wild-type copies of the region and a much longer amplicon from the copy carrying the docking platform. But when GFP^-^ cells were cloned from a population that had been challenged with an attB targeting plasmid, the same PCR reaction almost always failed to produce any amplicon other than the small wild-type product. We conclude that loss of the genomic region containing the docking site (see above) occurs at a much higher frequency than targeted substitution; selection of GFP^-^ cells was of little utility in isolating the correctly targeted products.
2. MTX selection of the transfected population efficiently selected for cells that carried the targeting construct. But when we used PCR directed at attB, attL, and attR sites, we found that the vast majority of clones made from this population retained attB sites, and only a very small fraction had the recombinant attL and attR sites generated by targeted integration(data not shown). We conclude that illegitimate recombination (in which the entire targeting plasmid is integrated occurs at much higher than targeted substitution.

In order to permit selection against cells in which the entire attB plasmid has been incorporated by illegitimate recombination, we modified our original attB vectors by adding a HS-TK transcription unit, whose product renders cells sensitive to ganciclivir (GCV); these modified vectors are shown in Figure 1, and were used in all of the remaining experiments described in this paper. Since targeted substitution incorporates only the region flanked by attB sites, GCV selection should kill cells in which the entire plasmid is incorporated and spare those in which only targeted replacement has occurred. Thus, treatment with MTX to kill cells in which the targeting plasmid has not been incorporated plus GCV to kill cells in which the targeting plasmid has been incorporated illegitimately should to enrich the population for cells with the intended targeted substitution only.

We began by targeting the two docking site lines with the empty attB vector B-DHFR-B-TK (Fig. 1). We selected with MTX, followed by MTX+GCV. MTX selection was complete within 2 weeks, as reported previously (Cherbas *et al.* 1994a). GCV selection was inefficient, as shown by the fact that control populations transfected with the attB vector in the absence of integrase displayed only a transient slowing of growth when treated with GCV. The MTX^R^ GCV^R^ population was then cloned in order to isolate a homogeneous population of cells which contained only the correctly targeted insert

We used PCR directed at attB, attL, and attR to distinguish between targeted insertions (which contain attL and attR, but no attB), illegitimate insertions (attB but no attL or attR), and a mixture of the two. The great majority of clones from populations selected with MTX and GCV and failing to express GFP contained both correct and illegitimate insertions (data not shown). The integrase expression plasmid was incorporated into the genome in only a very small minority of clones.

In a separate experiment we targeted B-DHFR-mCherry-B-TK to the same two docking site lines (Table 2). The results were very similar to those seen with the empty vector, but in this case we also assayed Cu^++^-inducible expression of mCherry in clones which contained only a correct insert by the attL/attR/attB test. Surprisingly, a significant number of clones failed to express any detectable mCherry; this was particularly true when a relatively high level of act-integrase was used in the transfection, and when the docking site Kc167-IPPI-66D was targeted. We speculate that following targeted insertion, a secondary integrase reaction involving cryptic attB and/or attP sites leads to a rearrangement of the DNA within the docking site; this notion is supported by the fact that these clones which showed correct formation of attL and attR sites and loss of GFP expression, and by the fact that loss of mCherry expression seems to be correlated with the amount of integrase expression plasmid used in the transfection, Further experiments will be required to determine the nature of the rearrangement, but we are encouraged that such secondary rearrangements seem to occur relatively rarely at low concentrations of act-integrase.

**Table 2.** Products of RMCE transfection in Kc167 docking site lines. Data refer to clonal lines derived from the procedure described above, targeting B-DHFR-Mt-mCherry-B-TK to the docking platforms in Kc167-PP-93E and Kc167-IPPI-66D. Transfections included 1 or 2 μg of act-integrase. All clones were MTX-resistant and had no detectable GFP. The table indicates the number of clones that were expanded and analyzed, and the fraction that passed the tests for correct integration only (attB absent, attL and attR present by PCR assay), stable integration of the integrase plasmid (integrase present by PCR assay), and expression of mCherry (FACS analysis of Cu^++^-treated cells). For each fraction, the denominator is the number of clones from the previous step that were tested.

| | IPPI-66D (1<br>$\mu\text{g}$<br>integrase) | IPPI-66D<br>(2 $\mu\text{g}$<br>integrase) | PP-93E (1<br>$\mu\text{g}$<br>integrase) | PP-93E (2<br>$\mu\text{g}$<br>integrase) |
| --- | --- | --- | --- | --- |
| total | 14 | 10 | 12 | 37 |
| passed<br>B/R/L test | 11/14 | 9/10 | 12/12 | 33/37 |
| integrase<br>retained | 1/11 | 0/10 | 0/12 | 2/35 |
| mCherry<br>expressed | 7/10 | 0/5 | 6/6 | 6/7 |

On the basis of the experiments just described, we recommend the following procedure for targeted replacement in our IPPI and PP docking site lines:

1. Co-transfect 1 ml of a docking site cell line, at about 3-5×10^6^ cells/ml, using Lipofectamine LTX with PLUS reagent (Life Technologies. For each transfection, use 0.5 μg act-φC31-integrase plus 2 μg of B-Mt-mCherry-DHFR-B-TK (8.0 kbp) or an equivalent molar concentration of a similar attB targeting plasmid.
2. After 2 days, transfer the cells to 5 ml of medium containing MTX (2×10^−7^ M final concentration). Change the medium every 4 days, retaining the MTX and diluting the cells as necessary to keep their concentration <1×10^7^/ml, until healthy MTX-resistant cells dominate the population; this process generally takes about 2 weeks.
3. Add GCV (20 μM final concentration) to the medium, and continue maintaining the cells in the presence of MTX and GCV, changing the medium every 4 days, until the growth rate is clearly depressed compared to a control culture containing only MTX; this process generally takes about 2-3 weeks.
4. Using FACS, clone GFP^-^ cells. When growing clones are clearly visible, use a fluorescence microscope to confirm the absence of GFP, and expand the GFP^-^ clones. In the case of Mt-mCherry targeting, we were able to get similar results with a much smaller background of GFP^+^ clones by treating the population for 24 prior to cloning with CuSO_4_, and then cloning GFP-mCherry^+^ cells.
5. Prepare DNA from candidate clones and use PCR to confirm the presence of attL and attR sites, the absence of attB sites, and the absence of the integrase plasmid.
6. Test candidate clones for expression of the targeting transgene (mCherry in our example) to eliminate clones with secondary rearrangements.

### Properties of clones carrying targeted insertions.

Stably transformed lines are routinely generated in *Drosophila* cell lines by illegitimate recombination between tandem arrays of transfected plasmids and random genomic sites (Moss 1985; Cherbas *et al.* 1994a). We expected that insertion of a single copy of a transgene into a docking site might give both increased stability of the transgene structure and more nearly normal chromatin structure and expression regulation than illegitimate insertion of tandem arrays of the same construct. For this reason, we compared the properties of cells containing either targeted or illegitimate insertions of an identical Mt-mCherry transcription unit. To maximize the homogeneity of the cell populations, we restricted ourselves to clonal lines. For illegitimate insertions, we used transfections that included the attB-Mt-mCherry donor plasmid but not a source of φC31 integrase; transformed cells were selected for resistance to MTX, and then cloned. For targeted insertions, we used the procedure described above. Examples of mCherry expression patterns are shown in Figure 3. For each line, a FACS histogram of mCherry expression is shown in untreated cells, and in cells treated for 20 hr with 1 mM CuSO_4_ to induce the expression of the Mt promoter. Autofluorescence was estimated from Kc167 cells (panel A), which do not contain a coding sequence for mCherry. The two targeted lines (B and C) have low background expression in the absence of Cu^++^ treatment, and a strong induction by Cu^++^; the two lines differ both in the intensity and the uniformity of mCherry expression. The photomicrograph in panel G illustrates both the variation in intensity of mCherry expression and the complete absence of nuclear GFP that is expected in targeted substitution. By contrast, clones with illegitimate arrays of Mt-mCherry (D, E, F, H) show a very broad range of background expression which is shifted to higher mCherry expression following Cu^++^ treatment, and retention of nuclear GFP expression; the 3 illegitimate clones shown in Figure 3 differ in the intensity of mCherry expression (both with and without Cu^++^ treatment) and the variation among cells in the population. In all cases, the targeted transformants show lower background expression, higher induction ratio, and lower variation among cells is lower than the illegitimate transformants.

## DISCUSSION

We report here a series of transformants made from Kc167 and from Sg4, each carrying a single copy of a docking site bearing 2 φC31 attP sites, designed for integrase-mediated cassette. In this communication we provide protocols for targeting transgenes to the docking sites, and describe vectors for preparation of attB-bounded constructs for this purpose.

### Distribution of P element insertion sites.

Data in this paper provide the first mappings of P element insertion sites in somatic cells. FlyBase release FB2015_03 includes mappings of 67,543 separate P element insertions into the germ line. Those sites are not randomly distributed; they show a preference for localized structural features (Liao *et al.* 2000) and an association with DNA replication origins (Spradling *et al.* 2011) and transcriptional activity (Fontanillas *et al.* 2007). P element “hot spots” are usually located in promoter regions (Spradling *et al.* 1995), but only 2% of promoters accounted for over 40% of P element insertions in a recent survey of over 18,000 independent transpositions.

Although our data for insertions into somatic cell lines is much too sparse for careful statistical comparison with those for germ-line insertion sites, even these restricted data are sufficient to show that the somatic insertions map preferentially in the vicinity of germ-line hot spots, despite large differences in the patterns of transcription and of replication origins (Cherbas *et al.* 2011; Eaton *et al.* 2011; Graveley *et al.* 2011; Brown *et al.* 2014). To illustrate this point, we note that most 2-kb segments of the genome have no known germ-line P element insertions; using the data from FlyBase, we surveyed 100 such segments, and found that 82 of them contained no mapped germ-line sites (data not shown). Yet 13 of the 17 insertions into Kc167 or Sg4 lay within 1 kb of at least 1 known germ-line site, and 8 of the 17 lay within 1 kb of at least 10 germ-line sites. Figure 4 shows a typical example of a docking site insertion which is clearly located in a hot spot for germ-line insertions.

**Figure 4.**
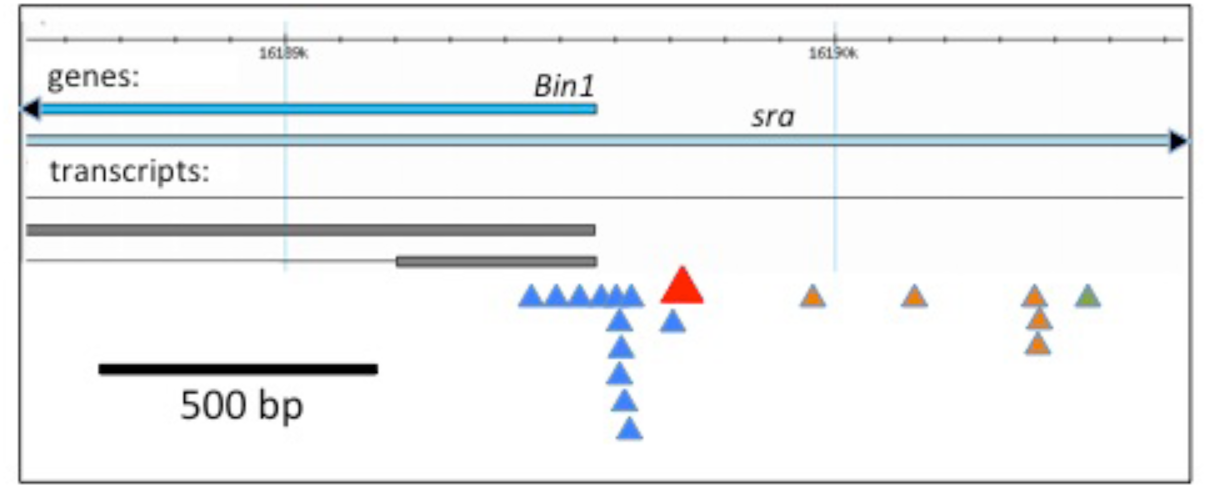
Insertion site for the docking platform in Kc167-PP-89B in a hot spot for P element insertions. The map is simplified from a FlyBase GBrowse view (DOS SANTOS *et al.* 2015) showing a 2.5 kb region of chromosome arm 3R; the insertion site for the P element docking platform transposon is indicated as a red arrowhead in the context of transposon insertion sites previously mapped in transformed flies. The region illustrated in this figure includes the promoter for the gene *Bin1*, and is entirely contained within an intron of the gene *sra*. Known transposon insertion sites in flies are shown as color-coded arrowheads: blue, P element; brown, PiggyBac; green, Minos.

### Targeting: problems in cell lines.

The efficiency of GCV selection as reported in mammalian cells is quite variable (Seibler *et al.* 1998; Converse *et al.* 2004; Chakraborty *et al.* 2013), with the strength of the promoter driving HS-TK probably playing an important role. A similar experiment in a *Drosophila* cell line, in which HS-TK was driven by an *Act5C*-GAL4 driver combined with a UAS-promoter gave efficient GCV selection (Manivannan *et al.* 2015), and we suspect that the difference in selection efficiency between the two results is that the amplification of TK expression produced by the GAL4/UAS system. We chose to use the *Act5C* promoter, a standard strong constitutive promoter in these cell lines, without the amplification conferred by the GAL4/UAS system, in order not to preclude other possible uses of GAL4/UAS in subsequent experiments using the targeted cells. The consequence of this decision is that GCV selection only provides enrichment for cells carrying the targeted substitution and no illegitimate insertions of the targeting plasmid, but in order to achieve a pure population, it is necessary to clone the cells and use PCR to identify the desired clones.

When targeted replacement is done in flies, illegitimate insertions do not occur, and scoring the loss of a *w*^+^ marker from docking site and the appearance of a *y*^+^ marker from the donor cassette was sufficient to give efficient identification of flies with the correct structure (Bateman *et al.* 2006).

### Targeting: Properties of the resulting lines.

Using 2 of the docking site lines, one with and one without insulator elements, we have characterized the expression of a Cu^++^-inducible Mt-mCherry marker inserted into the docking site. The properties of the targeted transgene are remarkably different from the properties of clones carrying the same transgene transformed by current methods in their stability, and in their expression properties.

Currently, stable transformation of *Drosophila* cell lines is usually done by introduction of the exogenous DNA, using any of a variety of techniques, and then selecting cells in which the DNA is incorporated into the genome. Incorporation into the genome occurs by illegitimate insertion, generally preceded by the formation of long arrays of the exogenous plasmid through homologous recombination. The insertion of these long arrays into the chromosome has been described extensively only for S2 cells transfected using calcium phosphosphate-DNA co-precipitates (Moss 1985; Cherbas *et al.* 1994a); more fragmentary data indicates that a similar process occurs in Kc167, though the length of the array may vary with the transfection procedure (Bourouis and Jarry 1983; Lee 1990). The products of these transformations are reasonably stable, as observed at the level of Southern blots (Moss 1985), but there is substantial variation from cell to cell. Such variation is to be expected in an uncloned population, but it occurs even in clonal populations ((Fehon *et al.* 1990; Lee 1990) and Figure 3), consistent with the tendency of tandem arrays undergo frequent deletions and expansions through homologous recombination. A single copy of a transgene inserted by targeted substitution would be expected to be much more stable and homogeneous; our measurements confirm this expectation (Figure 3). Insertion of a single copy of a transgene by P-element transposition (Segal *et al.* 1996) also produces clones with reasonably homogeneous expression (Figure 2); this procedure has been available for 20 years, but has been rarely used because it is much more cumbersome than illegitimate insertion. The targeted substitution procedure described in this paper is predicated on the insertion of single copies of a docking platform by P element transposition; once the docking site lines are isolated and characterized, constructs may be targeted to these docking sites. The targeting step requires more time and effort than illegitimate integration and can be recommended only for applications where increased homogeneity and improved transcriptional regulation of the targeted inserts can compensate for the extra work. As more of these targeted transformants become available, we expect that they may be particularly valuable as substrates for rapidly emerging CRISPR/Cas techniques for mutation and replacement (Bassett *et al.* 2014).

Our approach differs from that developed in the Simcox lab (Manivannan *et al.* 2015) in several respects, each conferring both advantages and disadvantages. First, and most important, we have chosen to start with existing, well-characterized cell lines rather than establishing new cell lines from existing fly stocks carrying well-characterized docking sites. This makes it possible to make use of the extensive data already available for the parental lines. Second, we have chosen to use attP sites (and therefore attB sites) in parallel rather than opposing orientation. This fixes the orientation of the resulting insertion. Third, we have used the Actin5C promoter to drive expression of constitutive markers, such as GFP, MTX, and GCV, rather than the much stronger combination of Act5C-GAL4 and a UAS promoter. This has the advantage of not interfering with the use of GAL4 drivers for experiments using the targeted line once it is established, but the disadvantage of making GCV selection inefficient, as described above.

## Acknowledgments

We are grateful to Jie Huang for technical assistance, to Amanda Simcox for helpful discussions and sharing data prior to publication, to Christiane Hassel and the IU Flow Cytometry Core Facility for experimental equipment and advice, to James Powers and the IUB Light Microscopy Imaging Center for help with fluorescence photomicrography, to Mimi Zolan for the use of her cesium source, to Greg Crouch and Amanda McKeen for performing the irradiations, and to Craig Pikaard and his laboratory for allowing us to use their ddPCR machine and for helpful advice. This work was supported by NIH grant 2P40OD010949-10A1.

## Abbreviations

act5C: Actin5C promoter
ddPCR: digital drop PCR
FCS: fetal calf serum
GCV: ganciclivir
HS-TK: Herpes simplex thymidine kinase
Mt: MtnA promoter
MTX: methotrexate
PCR: polymerase chain reaction

## References

Alekseyenko, A. A., J. W. Ho, S. Peng, M. Gelbart, M. Y. Tolstorukov et al., 2012 Sequence-specific targeting of dosage compensation in Drosophila favors an active chromatin context. PLoS Genet 8: e1002646.

Arndt-Jovin, D., 2006, pp., edited by L. Cherbas.

Barolo, S., L. A. Carver and J. W. Posakony, 2000 GFP and beta-galactosidase transformation vectors for promoter/enhancer analysis in Drosophila. Biotechniques 29: 726, 728, 730, 732.

Bassett, A. R., C. Tibbit, C. P. Ponting and J. L. Liu, 2014 Mutagenesis and homologous recombination in Drosophila cell lines using CRISPR/Cas9. Biol Open 3: 42–49.

Bateman, J. R., J. E. Johnson and M. N. Locke, 2012 Comparing enhancer action in cis and in trans. Genetics 191: 1143–1155.

Bateman, J. R., A. M. Lee and C. T. Wu, 2006 Site-specific transformation of Drosophila via phiC31 integrase-mediated cassette exchange. Genetics 173: 769–777.

Bateman, J. R., M. F. Palopoli, S. T. Dale, J. E. Stauffer, A. L. Shah et al., 2013 Captured segment exchange: a strategy for custom engineering large genomic regions in Drosophila melanogaster. Genetics 193: 421–430.

Bischof, J., R. K. Maeda, M. Hediger, F. Karch and K. Basler, 2007 An optimized transgenesis system for Drosophila using germ-line-specific phiC31 integrases. Proc Natl Acad Sci U S A 104: 3312–3317.

Bourouis, M., and B. Jarry, 1983 Vectors containing a prokaryotic dihydrofolate reductase gene transform Drosophila cells to methotrexate-resistance. EMBO J 2: 1099–1104.

Brown, J. B., N. Boley, R. Eisman, G. E. May, M. H. Stoiber et al., 2014 Diversity and dynamics of the Drosophila transcriptome. Nature 512: 393–399.

Bunch, T. A., Y. Grinblat and L. S. Goldstein, 1988 Characterization and use of the Drosophila metallothionein promoter in cultured Drosophila melanogaster cells. Nucleic Acids Res 16: 1043–1061.

Chakraborty, S., N. Christoforou, A. Fattahi, R. W. Herzog and K. W. Leong, 2013 A robust strategy for negative selection of Cre-loxP recombination-based excision of transgenes in induced pluripotent stem cells. PLoS One 8: e64342.

Cherbas, L., and P. Cherbas, 1997 “Parahomologous” gene targeting in Drosophila cells: an efficient, homology-dependent pathway of illegitimate recombination near a target site. Genetics 145: 349–358.

Cherbas, L., and P. Cherbas, 2007 Transformation of *Drosophila* cell lines: an alternative approach to exogenous protein expression, pp. 317–340 in Baculovirus Expression Protocols, edited by D. W. Murhammer. Humana Press.

Cherbas, L., and L. Gong, 2014 Cell lines. Methods 68: 74–81.

Cherbas, L., R. Moss and P. Cherbas, 1994a Transformation techniques for Drosophila cell lines. Methods Cell Biol 44: 161–179.

Cherbas, L., R. Moss and P. Cherbas, 1994b Transformation techniques for Drosophila cell lines. Methods in Cell Biology 44: 161–179.

Cherbas, L., A. Willingham, D. Zhang, L. Yang, Y. Zou et al., 2011 The transcriptional diversity of 25 Drosophila cell lines. Genome Res 21: 301–314.

Converse, A. D., L. R. Belur, J. L. Gori, G. Liu, F. Amaya et al., 2004 Counterselection and co-delivery of transposon and transposase functions for Sleeping Beauty-mediated transposition in cultured mammalian cells. Biosci Rep 24: 577–594.

DasGupta, R., A. Kaykas, R. T. Moon and N. Perrimon, 2005 Functional genomic analysis of the Wnt-wingless signaling pathway. Science 308: 826–833.

Dewey, R. A., G. Morrissey, C. M. Cowsill, D. Stone, F. Bolognani et al., 1999 Chronic brain inflammation and persistent herpes simplex virus 1 thymidine kinase expression in survivors of syngeneic glioma treated by adenovirus-mediated gene therapy: implications for clinical trials. Nat Med 5: 1256–1263.

dos Santos, G., A. J. Schroeder, J. L. Goodman, V. B. Strelets, M. A. Crosby et al., 2015 FlyBase: introduction of the Drosophila melanogaster Release 6 reference genome assembly and large-scale migration of genome annotations. Nucleic Acids Res 43: D690–697.

Eaton, M. L., J. A. Prinz, H. K. MacAlpine, G. Tretyakov, P. V. Kharchenko et al., 2011 Chromatin signatures of the Drosophila replication program. Genome Res 21: 164–174.

Echalier, G., and A. Ohanessian, 1969 Isolement, en cultures in vitro, de lignees cellulaires diploides de *Drosophila melanogaster*. C. r. hebd. Seanc. Acad. Sci., Paris D. Sci. Nat. 268: 1771–1773.

Ejsmont, R. K., and B. A. Hassan, 2014 The Little Fly that Could: Wizardry and Artistry of Drosophila Genomics. Genes (Basel) 5: 385–414.

Fehon, R. G., P. J. Kooh, I. Rebay, C. L. Regan, T. Xu et al., 1990 Molecular interactions between the protein products of the neurogenic loci Notch and Delta, two EGF-homologous genes in Drosophila. Cell 61: 523–534.

Fish, M. P., A. C. Groth, M. P. Calos and R. Nusse, 2007 Creating transgenic Drosophila by microinjecting the site-specific phiC31 integrase mRNA and a transgene-containing donor plasmid. Nat Protoc 2: 2325–2331.

Fontanillas, P., D. L. Hartl and M. Reuter, 2007 Genome organization and gene expression shape the transposable element distribution in the Drosophila melanogaster euchromatin. PLoS Genet 3: e210.

Fujioka, M., G. L. Yusibova, J. Zhou and J. B. Jaynes, 2008 The DNA-binding Polycomb-group protein Pleiohomeotic maintains both active and repressed transcriptional states through a single site. Development 135: 4131–4139.

Gloor, G. B., C. R. Preston, D. M. Johnson-Schlitz, N. A. Nassif, R. W. Phillis et al., 1993 Type I repressors of P element mobility. Genetics 135: 81–95.

Goetze, S., A. Baer, S. Winkelmann, K. Nehlsen, J. Seibler et al., 2005 Performance of genomic bordering elements at predefined genomic loci. Mol Cell Biol 25: 2260–2272.

Graveley, B. R., A. N. Brooks, J. W. Carlson, M. O. Duff, J. M. Landolin et al., 2011 The developmental transcriptome of *Drosophila melanogaster*. Nature 471: 473–479.

Groth, A. C., M. Fish, R. Nusse and M. P. Calos, 2004 Construction of transgenic Drosophila by using the site-specific integrase from phage phiC31. Genetics 166: 1775–1782.

Hindson, B. J., K. D. Ness, D. A. Masquelier, P. Belgrader, N. J. Heredia et al., 2011 High-throughput droplet digital PCR system for absolute quantitation of DNA copy number. Anal Chem 83: 8604–8610.

Huang, J., W. Zhou, W. Dong and Y. Hong, 2009a Targeted engineering of the Drosophila genome. Fly (Austin) 3: 274–277.

Huang, J., W. Zhou, W. Dong, A. M. Watson and Y. Hong, 2009b From the Cover: Directed, efficient, and versatile modifications of the Drosophila genome by genomic engineering. Proc Natl Acad Sci U S A 106: 8284–8289.

Karpova, N., Y. Bobinnec, S. Fouix, P. Huitorel and A. Debec, 2006 Jupiter, a new *Drosophila* protein associated with microtubules. Cell Motil Cytoskeleton 63: 301–312.

Koppen, T., A. Weckmann, S. Muller, S. Staubach, W. Bloch et al., 2011 Proteomics analyses of microvesicles released by Drosophila Kc167 and S2 cells. Proteomics 11: 4397–4410.

Lau, N. C., N. Robine, R. Martin, W. J. Chung, Y. Niki et al., 2009 Abundant primary piRNAs, endo-siRNAs, and microRNAs in a Drosophila ovary cell line. Genome Res 19: 1776–1785.

Lee, H., C. McManus, D. Y. Cho, M. Eaton, F. Renda et al., 2014a DNA copy number evolution in Drosophila cell lines. Genome Biol 15: R70.

Lee, H., C. J. McManus, D. Y. Cho, M. Eaton, F. Renda et al., 2014b DNA copy number evolution in Drosophila cell lines. Genome Biol 15: R70.

Lee, K., 1990 The Identification and Characterization of Ecdysone Response Elements, pp. 263 in Department of Biology. Indiana University.

Liao, G. C., E. J. Rehm and G. M. Rubin, 2000 Insertion site preferences of the P transposable element in Drosophila melanogaster. Proc Natl Acad Sci U S A 97: 3347–3351.

Liu, T., D. Sims and B. Baum, 2009 Parallel RNAi screens across different cell lines identify generic and cell type-specific regulators of actin organization and cell morphology. Genome Biol 10: R26.

Manivannan, S. N., T. L. Jacobsen, P. Lyon, B. Selvaraj, P. Halpin et al., 2015 Targeted integration of single-copy transgenes in *Drosophila melanogaster* tissue culture cells using recombination-mediated cassette exchange, pp., submitted to Genetics.

Mirkin, E. V., F. S. Chang and N. Kleckner, 2014 Protein-mediated chromosome pairing of repetitive arrays. J Mol Biol 426: 550–557.

Monroe, T. J., M. C. Muhlmann-Diaz, M. J. Kovach, J. O. Carlson, J. S. Bedford et al., 1992 Stable transformation of a mosquito cell line results in extraordinarily high copy numbers of the plasmid. Proc Natl Acad Sci U S A 89: 5725–5729.

Moss, R. E., 1985 Analysis of a transformation system for *Drosophila* tissue culture cells, pp. in Department of Biology. Harvard University.

Neumuller, R. A., F. Wirtz-Peitz, S. Lee, Y. Kwon, M. Buckner et al., 2012 Stringent analysis of gene function and protein-protein interactions using fluorescently tagged genes. Genetics 190: 931–940.

Potter, C. J., and L. Luo, 2010 Splinkerette PCR for mapping transposable elements in Drosophila. PLoS One 5: e10168.

Riddle, N. C., Y. L. Jung, T. Gu, A. A. Alekseyenko, D. Asker et al., 2012 Enrichment of HP1a on Drosophila chromosome 4 genes creates an alternate chromatin structure critical for regulation in this heterochromatic domain. PLoS Genet 8: e1002954.

Riddle, N. C., A. Minoda, P. V. Kharchenko, A. A. Alekseyenko, Y. B. Schwartz et al., 2011 Plasticity in patterns of histone modifications and chromosomal proteins in Drosophila heterochromatin. Genome Res 21: 147–163.

Robb, J. A., 1969 Maintenance of imaginal discs of *Drosophila melanogaster* in chemically defined media. J. Cell Biology 41: 876–885.

Rosser, J. M., and W. An, 2010 Repeat-induced gene silencing of L1 transgenes is correlated with differential promoter methylation. Gene 456: 15–23.

Rubin, G. M., and A. C. Spradling, 1983a Vectors for P element-mediated gene transfer in Drosophila. Nucleic Acids Res 11: 6341–6351.

Rubin, G. M., and A. C. Spradling, 1983b Vectors for P element-mediated gene transfer in *Drosophila*. Nucleic Acids Research 11: 6341–6351.

Schaaf, C. A., Z. Misulovin, G. Sahota, A. M. Siddiqui, Y. B. Schwartz et al., 2009 Regulation of the Drosophila Enhancer of split and invected-engrailed gene complexes by sister chromatid cohesion proteins. PLoS One 4: e6202.

Schneider, I., 1972 Cell lines derived from late embryonic stages of *Drosophila melanogaster*. Journal of Embryology and Experimental Morphology 27: 353– 365.

Schwartz, Y. B., T. G. Kahn, P. Stenberg, K. Ohno, R. Bourgon et al., 2010 Alternative epigenetic chromatin states of polycomb target genes. PLoS Genet 6: e1000805.

Segal, D., L. Cherbas and P. Cherbas, 1996 Genetic transformation of *Drosophila* cells in culture by P element-mediated transposition. Somat Cell Mol Genet 22: 159–165.

Seibler, J., D. Schubeler, S. Fiering, M. Groudine and J. Bode, 1998 DNA cassette exchange in ES cells mediated by Flp recombinase: an efficient strategy for repeated modification of tagged loci by marker-free constructs. Biochemistry 37: 6229–6234.

Spradling, A. C., H. J. Bellen and R. A. Hoskins, 2011 Drosophila P elements preferentially transpose to replication origins. Proc Natl Acad Sci U S A 108: 15948–15953.

Spradling, A. C., D. M. Stern, I. Kiss, J. Roote, T. Laverty et al., 1995 Gene disruptions using P transposable elements: an integral component of the Drosophila genome project. Proc Natl Acad Sci U S A 92: 10824–10830.

Sun, F. F., J. E. Johnson, M. P. Zeidler and J. R. Bateman, 2012 Simplified Insertion of Transgenes Onto Balancer Chromosomes via Recombinase-Mediated Cassette Exchange. G3 (Bethesda) 2: 551–553.

Vatolina, T. Y., L. V. Boldyreva, O. V. Demakova, S. A. Demakov, E. B. Kokoza et al., 2011 Identical functional organization of nonpolytene and polytene chromosomes in Drosophila melanogaster. PLoS One 6: e25960.

Venken, K. J., and H. J. Bellen, 2012 Genome-wide manipulations of Drosophila melanogaster with transposons, Flp recombinase, and PhiC31 integrase. Methods Mol Biol 859: 203–228.

Venken, K. J., Y. He, R. A. Hoskins and H. J. Bellen, 2006 P[acman]: a BAC transgenic platform for targeted insertion of large DNA fragments in D. melanogaster. Science 314: 1747–1751.

Venken, K. J., E. Popodi, S. L. Holtzman, K. L. Schulze, S. Park et al., 2010 A molecularly defined duplication set for the X chromosome of Drosophila melanogaster. Genetics 186: 1111–1125.

Wen, J., J. Mohammed, D. Bortolamiol-Becet, H. Tsai, N. Robine et al., 2014 Diversity of miRNAs, siRNAs, and piRNAs across 25 Drosophila cell lines. Genome Res 24: 1236–1250.

Weng, R., Y. W. Chen, N. Bushati, A. Cliffe and S. M. Cohen, 2009 Recombinase-mediated cassette exchange provides a versatile platform for gene targeting: knockout of miR-31b. Genetics 183: 399–402.

Williams, B. R., J. R. Bateman, N. D. Novikov and C. T. Wu, 2007 Disruption of topoisomerase II perturbs pairing in drosophila cell culture. Genetics 177: 31–46.

Wurtele, H., K. C. Little and P. Chartrand, 2003 Illegitimate DNA integration in mammalian cells. Gene Ther 10: 1791–1799.

Zhang, X., W. H. Koolhaas and F. Schnorrer, 2014 A Versatile Two-Step CRISPR-and RMCE-Based Strategy for Efficient Genome Engineering in Drosophila. G3 (Bethesda).

Zurovec, M., T. Dolezal, M. Gazi, E. Pavlova and P. J. Bryant, 2002 Adenosine deaminase-related growth factors stimulate cell proliferation in Drosophila by depleting extracellular adenosine. Proc Natl Acad Sci U S A 99: 4403–4408.

